# Relationships between mediolateral step modulation and clinical balance measures in people with chronic stroke

**DOI:** 10.1101/2022.04.26.489530

**Authors:** Keith E. Howard, Nicholas K. Reimold, Heather L. Knight, Aaron E. Embry, Holly A. Knapp, Alexa A. Agne, Camden J. Jacobs, Jesse C. Dean

## Abstract

**Background:** Many people with chronic stroke (PwCS) exhibit walking balance deficits linked to increased fall risk and decreased balance confidence. One potential contributor to these balance deficits is a decreased ability to modulate mediolateral stepping behavior based on pelvis motion. This behavior, hereby termed mediolateral step modulation, is thought to be an important balance strategy but can be disrupted in PwCS.

**Research Question:** Are biomechanical metrics of mediolateral step modulation related to common clinical balance measures among PwCS?

**Methods:** In this cross-sectional study, 93 PwCS walked on a treadmill at their self-selected speed for 3-minutes. We quantified mediolateral step modulation for both paretic and non-paretic steps by calculating partial correlations between mediolateral pelvis displacement at the start of each step and step width (ρSW), mediolateral foot placement relative to the pelvis (ρFP), and final mediolateral location of the pelvis (ρPD) at the end of the step. We also assessed several common clinical balance measures (Functional Gait Assessment [FGA], Activities-specific Balance Confidence scale [ABC], self-reported fear of falling and fall history). We performed Spearman correlations to relate each biomechanical metric of step modulation to FGA and ABC scores. We performed Wilcoxon rank sum tests to compare each biomechanical metric between individuals with and without a fear of falling and a history of falls.

**Results:** Only ρFP for paretic steps was significantly related to all four clinical balance measures; higher paretic ρFP values tended to be observed in participants with higher FGA scores, with higher ABC scores, without a fear of falling and without a history of falls. However, the strength of each of these relationships was only weak to moderate.

**Significance:** While the present results do not provide insight into causality, they justify future work investigating whether interventions designed to increase ρFP can improve clinical measures of post-stroke balance in parallel.

## Introduction

Deficits in walking balance, defined here as a reduced ability to independently walk in one’s environment without falling, can limit function in many people with chronic stroke (PwCS). Supporting this idea, walking is the most common activity at the time of a fall in this population [1]. Many PwCS report a fear of falling [2] and reduced walking balance self-efficacy [3], which is linked to decreased community participation and quality of life [3,4]. Likely as a result, balance improvements are a primary rehabilitation goal for many patients [5].

While no existing measure can quantify an individual’s ability to walk in all potential environments without falling, numerous methods have been developed to assess walking balance. Commonly used clinical balance measures include the Functional Gait Assessment (FGA) to evaluate performance of walking tasks with a range of difficulty [6], the Activities-specific Balance Confidence scale (ABC) to quantify balance self-efficacy [7], and asking if a patient is afraid of falling [2]. While each of these measures has been linked to functional mobility [8,9] and may be used in clinical decision making, they do not provide insight into the mechanical underpinnings of post-stroke walking imbalance. An alternative approach is to quantify aspects of walking biomechanics that may indicate balance deficits. For example, relative to age-matched controls, PwCS tend to walk with slower speeds, shorter and wider steps, and more variable step lengths, times, widths, and margins of stability [10–13].

The present study focuses on mediolateral walking balance, which model simulations [14] and human experiments [15] suggest is more challenging to control than anteroposterior balance. Many post-stroke falls (∼31-38%) occur sideways toward the paretic side in both the chronic [16] and subacute [17] phases, supporting the clinical importance of mediolateral balance. The importance of mediolateral balance is further supported by the prospective prediction of post-stroke falls based on mediolateral motion of the pelvis during walking (e.g., lower mediolateral dynamic stability [18]; decreased mediolateral displacement [19]).

One strategy that plays an important role in mediolateral walking balance involves modulating the mediolateral motion of the body during a step based on the dynamic state of the pelvis, hereby termed mediolateral step modulation. In neurologically intact controls, step width varies with pelvis dynamics; steps tend to be wider when the pelvis has a larger mediolateral displacement away from the stance foot [20]. This relationship can be attributed to a combination of active control of swing leg position (influencing lateral foot placement relative to the pelvis [21]), and inverted pendulum dynamics of the stance leg and trunk (influencing the mediolateral position of the pelvis relative to the stance foot at the end of the step). Among PwCS, the relationship between mediolateral pelvis displacement and step width is weaker for steps taken with the paretic leg than the non-paretic leg [22], suggesting a disruption in the strategy used to achieve mediolateral step modulation. However, it is unclear whether this altered biomechanical behavior is related to common measures of walking balance.

The purpose of this exploratory, cross-sectional study was to investigate the potential link between mediolateral step modulation and common clinical balance measures among PwCS. Specifically, we quantified the relationship between mediolateral pelvis displacement at the start of each step and the body’s mediolateral configuration at the end of the step (i.e., step width, mediolateral foot placement relative to the pelvis, and mediolateral pelvis displacement relative to the stance foot) for both paretic and non-paretic steps. We then assessed the extent to which each of these biomechanical gait metrics was related to FGA score, ABC score, fear of falling, and self-reported fall history. The presence of significant links between biomechanical metrics and clinical balance measures could provide justification for interventions targeting the observed biomechanical behaviors.

## Methods

### Participants

93 PwCS participated in this study, which used data collected in a screening session for a separate clinical trial (NCT02960439). Inclusion criteria were: experience of a stroke ≥ 6 months before enrollment; age ≥ 21 years; ability to walk independently on a treadmill (e.g., without use of a cane, walker, or external assistance). Exclusion criteria were: resting heart rate above 110 beats/min; resting blood pressure above 200/110 mm Hg; history of congestive heart failure, unstable cardiac arrhythmias, hypertrophic cardiomyopathy, severe aortic stenosis, angina or dyspnea; preexisting neurological disorders or dementia; history of major head trauma; severe visual impairment; history of deep vein thrombosis or pulmonary embolism within 6 months; uncontrolled diabetes with recent weight loss, diabetic coma, or frequent insulin reactions; orthopedic injuries or conditions with the potential to alter the ability to adjust foot placement while walking. Participants were permitted to wear ankle foot orthoses if used during community walking.

Summary demographic and functional characteristics are presented in Table 1, with individual-level data available in Appendix A. The sample size for this study was based on the recruitment needs of the clinical trial. However, based on our goal of exploring associations between biomechanical and clinical measures, we surpassed the 84 participants needed to detect a moderate correlation (r=0.3) with 80% power and an alpha value of 0.05. All participants provided written informed consent, signing a form approved by the Medical University of South Carolina Institutional Review Board.

**Table 1.**
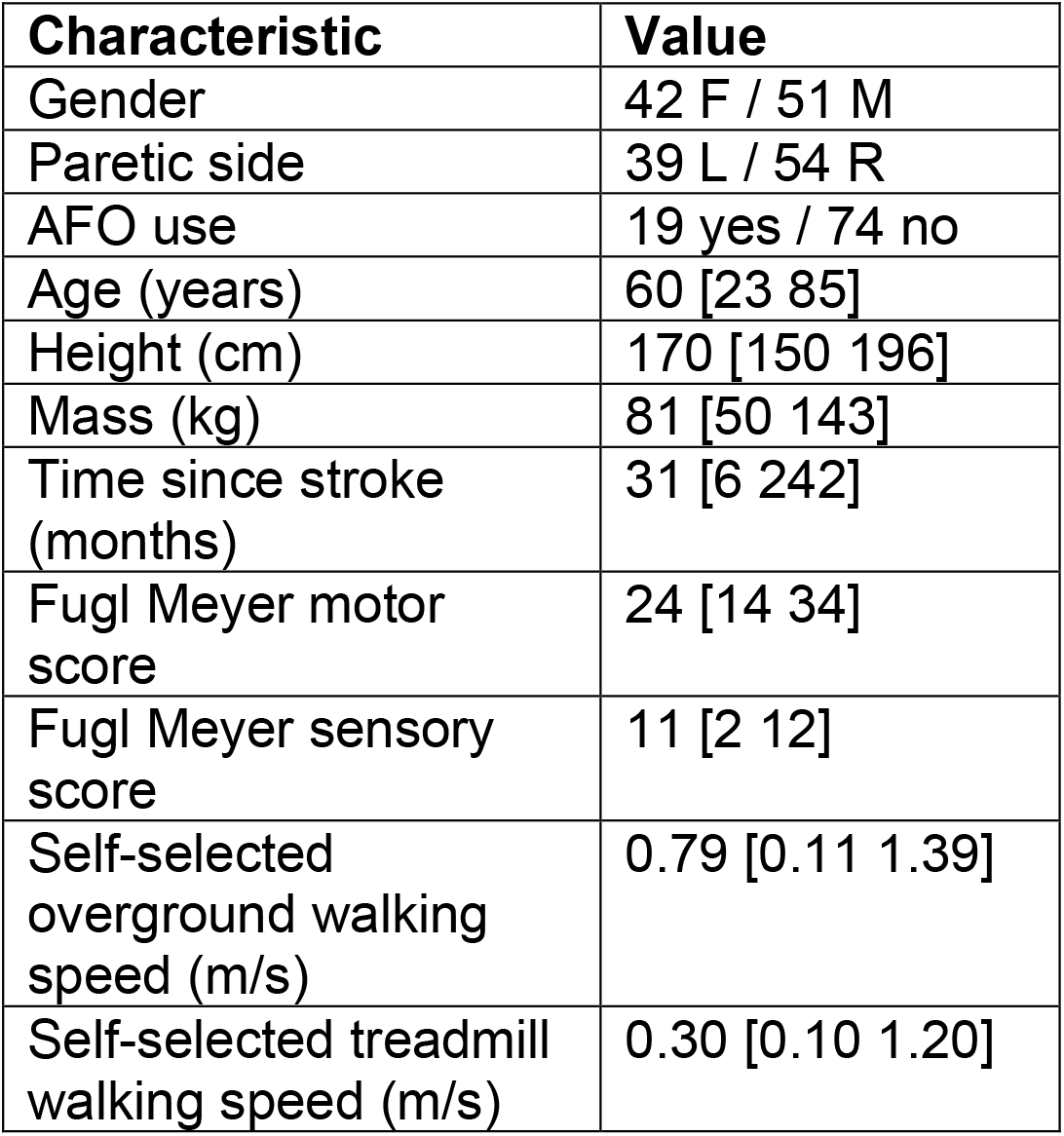
Summary participant information. For numerical measures, we present the group median and the group range in brackets.

| Characteristic | Value |
| --- | --- |
| Gender | 42 F / 51 M |
| Paretic side | 39 L / 54 R |
| AFO use | 19 yes / 74 no |
| Age (years) | 60 [23 85] |
| Height (cm) | 170 [150 196] |
| Mass (kg) | 81 [50 143] |
| Time since stroke (months) | 31 [6 242] |
| Fugl Meyer motor score | 24 [14 34] |
| Fugl Meyer sensory score | 11 [2 12] |
| Self-selected overground walking speed (m/s) | 0.79 [0.11 1.39] |
| Self-selected treadmill walking speed (m/s) | 0.30 [0.10 1.20] |

### Experimental Procedures – Clinical Testing

We calculated self-selected overground walking speed from the time required for participants to traverse the middle 6-meters of a 10-meter path, averaged across five passes. We quantified lower extremity Fugl-Meyer motor and sensory scores as measures of sensorimotor function, FGA scores to assess walking balance, ABC scores to quantify balance self-efficacy, and asked participants “Do you have a fear of falling?” to assess fear of falling [23]. We assessed self-reported fall history by asking participants “Have you had any fall in which you lost your balance and landed on a lower level in the past year?” [24]. All clinical testing procedures were performed by a licensed and trained physical therapist or physical therapist assistant.

### Experimental Procedures – Treadmill Walking

Biomechanical metrics were calculated from data collected while participants walked on a treadmill flush with the floor (Bertec; Columbus, OH; walking surface area of 173 cm by 63 cm). Participants did not hold on to a handrail, but wore a non-bodyweight support harness attached to an overhead rail for safety in case of a loss of balance.

We first identified each participant’s self-selected speed, defined for participants as the speed they used to walk around the house or a store. The treadmill started at 0.1 m/s and was increased in increments of 0.05 m/s until participants reported that the treadmill speed was faster than their self-selected speed, at which point the speed was reduced by 0.05 m/s. Participants walked for 3-minutes at the self-selected speed but were permitted to take breaks if they reported being unable to continue or were unable to maintain their position on the treadmill. Across participants, a maximum of three trials was required to complete the 3-minutes of walking.

### Data Collection and Step Identification

Active LED markers (PhaseSpace; San Leandro, CA) were placed on participants, with the present analyses focused on markers on the pelvis and bilateral heels. Marker position was measured at 120 Hz. In general, steps were defined to start when the foot’s velocity changed from posterior to anterior and to end when the foot’s velocity changed from anterior to posterior [25]. However, initial analyses revealed unusual foot velocity patterns in a subset of participants, motivating the use of a more complex method to avoid the detection of false gait events (detailed in Appendix B). Across participants, a median of 125 steps per leg (range = 62-181) were included in our analyses.

### Calculation of Biomechanical Metrics

At the start of each step, we calculated mediolateral pelvis displacement relative to the stance heel and mediolateral pelvis velocity, defining the positive direction as toward the swing leg. For each step, we calculated step width as the mediolateral displacement between the stance heel at the start of the step and the swing heel at the end of the step. Following prior methods used to quantify step width modulation [22,26,27], we calculated the partial correlation between step width and mediolateral pelvis displacement at the start of the step (ρSW), accounting for mediolateral pelvis velocity. Our use of partial correlations rather than R^2^ values is due to the presence of negative partial correlation values in some PwCS, which can artificially inflate R^2^ values [22]. Our focus on pelvis displacement is justified by the stronger link between step width and pelvis displacement than pelvis velocity [28], as well as our prior finding of significant post-stroke asymmetries only in the relationship between step width and pelvis displacement [22].

Step width modulation can be achieved by varying either the mediolateral placement of the swing foot relative to the pelvis (hereby termed *foot placement*) or the mediolateral displacement of the pelvis relative to the stance foot at the end of the step (hereby termed *final pelvis displacement*) (Fig. 1). Our prior work suggests that these two sources of step width modulation can vary independently [26]. Therefore, we calculated the partial correlation between foot placement and mediolateral pelvis displacement at the start of the step (ρFP), accounting for mediolateral pelvis velocity.

**Figure 1.**
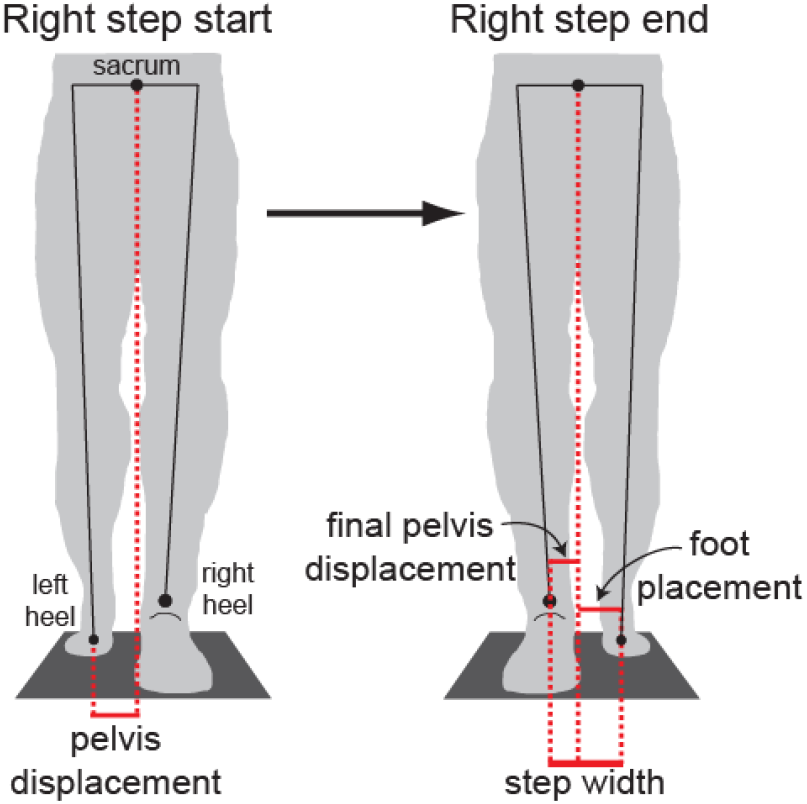
Schematic illustration of mediolateral step measurements, shown for a right step and adapted from [26]. At the start of each step (left illustration), we calculated mediolateral pelvis displacement between the sacrum and stance heel. At the end of the step (right illustration), we calculated final mediolateral pelvis displacement, mediolateral foot placement between the sacrum and swing heel, and step width. Step width was the sum of final pelvis displacement and foot placement.

This metric quantifies the extent to which the swing foot position is modulated based on pelvis displacement. We also calculated the partial correlation between final pelvis displacement and mediolateral pelvis displacement at the start of the step (ρPD), again accounting for mediolateral pelvis velocity (ρPD). This metric quantifies the extent to which the position of the pelvis at the end of a step can be attributed to its position at the start of the step.

While the present analyses focused on pelvis dynamics at the start of the step in order to focus on within-step adjustments in body configuration, an alternative approach quantifies the relationship between step width and pelvis dynamics at the end of the step [29]. For comparison, such analyses are included in Appendix C.

### Statistical Analyses

We explored potential relationships between our biomechanical metrics of mediolateral step modulation (ρSW, ρFP, and ρPD for both paretic and non-paretic steps) and several clinical balance measures. First, we used Spearman correlations to relate each of our biomechanical metrics to FGA score and ABC score. Correlation magnitudes of less than 0.3 are often interpreted as weak, between 0.3 and 0.7 as moderate, and greater than 0.7 as strong [30]. We used Wilcoxon rank sum tests to compare each of our biomechanical metrics between individuals with and without a fear of falling, as well as between individuals with and without a self-reported fall in the previous year. Effect size was quantified using Cliff’s delta, for which magnitudes less than 0.28 can be interpreted as weak, between 0.28 and 0.43 as moderate, and greater than 0.43 as strong [31]. For all comparisons, an alpha value of 0.05 was considered significant. Given the exploratory nature of our analyses, we did not correct for multiple comparisons.

## Results

Individual participant data points for all comparisons between biomechanical gait metrics and clinical balance measures are illustrated in Figure 2, along with the corresponding statistical test results and effect sizes. For all observed significant effects, the effect size magnitudes ranged from weak (0.22 for Spearman correlation and 0.26 for Cliff’s delta) to moderate (0.43 for Spearman correlation and 0.37 for Cliff’s delta). Individual participant values are available for all measures in Appendix A.

**Figure 2.**
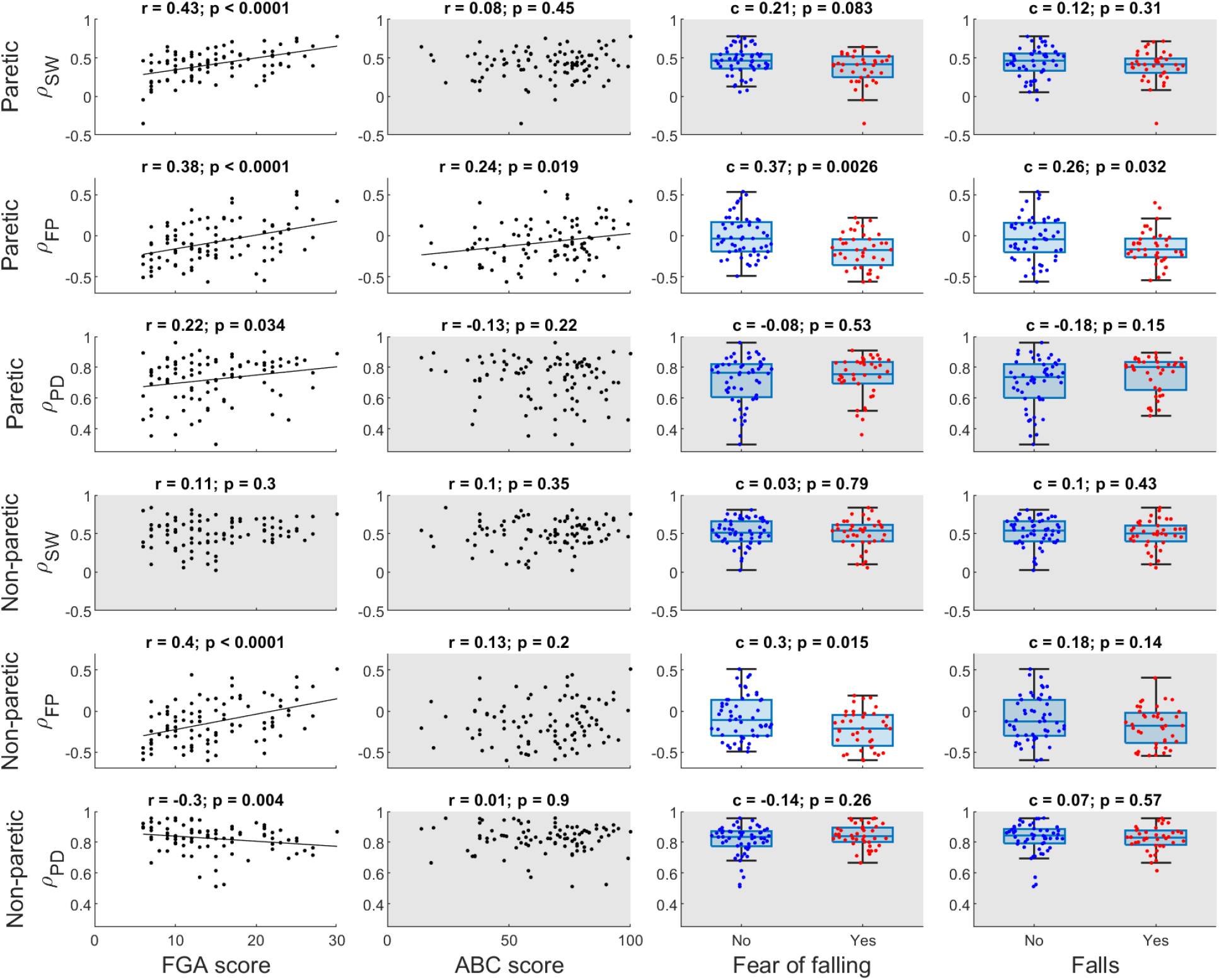
Comparison between biomechanical gait metrics (rows) and clinical balance measures (columns). For all panels, dots indicate individual participant values. Biomechanical gait metrics for paretic steps are plotted in the top three rows, and metrics for non-paretic steps are in the bottom three rows. In the first column, all six biomechanical gait metrics are plotted relative to FGA score. The effect size (r = Spearman correlation) and p-value are indicated at the top of each panel; panels without a significant relationship are shaded gray. In the case of a significant correlation, the best linear fit is shown for illustrative purposes. The second column follows the same structure for comparisons with ABC score. The third column presents box plots comparing biomechanical gait metrics between participants without and with a fear of falling. The central horizontal line indicates the median, the top and bottom edges of the box indicate the upper and lower quartile, and the whiskers indicate values 1.5 interquartile ranges above or below the corresponding quartile value (often used to identify outliers). The effect size (c = Cliff’s delta) and p-value are indicated at the top of each panel, which is shaded gray if no significant relationship was detected. The fourth column follows the same structure for comparisons between participants with and without a self-reported fall in the prior year.

Of the biomechanical gait metrics, only the partial correlation between pelvis displacement and mediolateral paretic foot placement (paretic ρFP) was significantly related to all four clinical balance measures. Specifically, paretic ρFP was higher for individuals with higher FGA scores, higher ABC scores, no fear of falling, and no self-reported falls.

For the remaining biomechanical gait metrics, both paretic ρSW and paretic ρPD were positively correlated with FGA score, but not significantly related to the other clinical balance measures. Non-paretic ρSW was not significantly related to any clinical balance measure. Non-paretic ρFP was positively correlated with FGA score and was higher for individuals without a fear of falling, but not significantly related to ABC score or self-reported falls. Finally, non-paretic ρPD was negatively correlated with FGA score, but not significantly related to the other clinical balance measures.

## Discussion

Our exploratory analyses revealed preliminary evidence for a link between altered mediolateral step modulation and balance in PwCS. Most notably, reduced modulation of mediolateral paretic foot placement was linked with poorer balance performance and confidence. However, all observed relationships were weak to moderate, indicating that the role of factors other than step modulation warrant investigation.

Modulation of mediolateral paretic foot placement (paretic ρFP) was significantly related to all clinical balance measures, albeit only weakly or moderately. Individuals with a stronger relationship between pelvis displacement and paretic foot placement tended to exhibit better balance and confidence. These results are consistent with a role of foot placement in walking balance [20], and suggest that control of where the swing foot lands is related to balance performance and self-efficacy in PwCS, both of which can influence real-world behavior [8]. The relatively weak relationships are perhaps not surprising, as the ability to adjust foot placement is only one component of the tasks assessed in the FGA, and many other factors surely contribute to balance confidence and the likelihood of real-world falls.

The other investigated biomechanical gait metrics were not consistently related to clinical balance measures, although a few significant relationships were observed. Greater modulation of mediolateral non-paretic foot placement (non-paretic ρFP) was linked with higher FGA scores and less fear of falling, likely because inaccurate placement of the non-paretic leg could influence walking balance similarly to inaccurate placement of the paretic leg. The significant relationships between modulation of final mediolateral pelvis displacement and FGA score were more complex. Higher FGA scores were weakly linked with greater pelvis displacement modulation during paretic steps (paretic ρPD). While speculative, this may be due to individuals with better balance allowing their stance non-paretic leg to act more like an inverted pendulum, with the mass of the body accelerating toward the paretic side. This behavior would subsequently require controlled weight transfer onto the paretic leg, which is limited in many PwCS with walking and balance deficits [32]. In contrast, *lower* FGA scores were linked with greater pelvis displacement modulation during non-paretic steps (non-paretic ρPD). Perhaps this is due to individuals with poor balance adopting walking strategies in which the paretic leg is placed far laterally [22], changing the initial mechanical conditions of the subsequent non-paretic step and allowing the stance paretic leg to act as an inverted pendulum and rapidly accelerate the mass of the body back toward their stronger non-paretic leg. Our simple measures of pelvis modulation are unable to disentangle these potential balance strategies, which would likely require more in-depth measures of muscle activity. Finally, greater modulation of paretic step width (paretic ρSW) was linked with higher FGA scores, paralleling the previously described result with mediolateral paretic foot placement.

A major limitation of this work was the lack of causal insight into relationships between biomechanical gait metrics and clinical balance measures. It is possible that a reduced ability to modulate paretic foot placement causes PwCS to perform more poorly at challenging balance tasks, and have lower balance self-efficacy, more fear of falling, and an increased risk of falls. Alternatively, individuals with more severe post-stroke deficits may simply be more likely to exhibit altered function during both treadmill walking and more functional tasks. Future experiments could gain causal insight by manipulating biomechanical gait metrics and determining whether clinical balance measures change in parallel. For example, we have demonstrated that targeted perturbations can influence step width modulation in both neurologically-intact controls [33] and PwCS [27], with a primary effect on ρFP [26]. It is presently unclear whether an intervention targeting increased paretic ρFP would produce parallel improvements in clinical balance measures.

A further limitation was our focus on a single biomechanical strategy for ensuring walking balance – mediolateral step modulation. While this strategy allows large base of support adjustments, it cannot influence body dynamics *within* a step. A faster alternative is a “center of pressure (CoP) shift strategy”, in which ankle musculature shifts the CoP location medially or laterally within the boundaries of the stance foot [34,35]. A “push-off strategy” in which the ankle musculature modulates trail leg propulsion [34], may also help ensure controlled mediolateral weight shift. Future work should investigate the use of these strategies in PwCS.

Additional limitations include our focus on unperturbed walking, which may not predict the ability to recover from perturbations [36]. Our requirement that participants are able to walk without a cane or walker restricted the study population to relatively high functioning individuals. Also, our results were derived from treadmill walking rather than overground walking. While this approach was necessary to feasibly measure many steps without risking participant fatigue, walking patterns can differ between these two contexts [37], most notably in the form of substantially lower speeds on a treadmill (here, median values of 0.30 m/s on a treadmill and 0.79 m/s overground). Similarly, the use of a harness was necessary to ensure safety, but may have emboldened participants and thus altered their gait pattern. Finally, falls were self-reported and retrospective, a method that can yield unreliable results [38]. Despite these limitations, the information gained from this type of experiment has value. Gait characteristics measured during steady-state treadmill walking have recently been found to be more promising predictors of prospective falls in PwCS than either clinical assessments or perturbation responses [18,39], possibly because post-stroke balance deficits are often due to internal movement errors [40] rather than altered reactions to external forces.

In conclusion, several biomechanical metrics of mediolateral step modulation were significantly linked with common clinical balance measures among PwCS. The most consistent links were observed with paretic ρFP, a metric of the extent to which paretic foot placement is modulated based on the dynamic state of the pelvis. However, the observed relationships were only weak to moderate, and do not provide insight into causality. Future work should investigate whether changes in foot placement modulation are accompanied by changes in clinical measures of balance performance.

## Acknowledgements

The views expressed by the authors are their own and do not necessarily reflect the official policy of the VA or the U.S. Government.

## Data statement

The data described in this manuscript is available in Appendix A.

