## Appendix B for "Relationships between mediolateral step modulation and clinical balance measures in people with chronic stroke"

To assess stepping behavior, we have quantified the partial correlation between mediolateral pelvis displacement and step width in several prior studies involving both neurologically-intact controls and people with chronic stroke (PwCS) [1–6]. In these studies, we defined the start and end of steps based on the anteroposterior velocities of markers placed on the heels, low-pass filtered at 10 Hz. Starting at the beginning of collected data, we identified the first timepoint at which the velocity of the left heel changed from negative to positive; this was defined at the start of the first left step. Moving forward in time, we identified the next timepoint at which the velocity of the contralateral right heel changed from negative to positive; this was defined as the end of the left step. This process then continued through the remainder of the trial and was repeated for right steps. Our definition of the end of the step was chosen because we assumed that the ipsilateral foot must be securely on the ground for the contralateral heel to begin moving forward, and that any lateral motion of the ipsilateral foot would be minimal once it contacted the ground.

The present study included a larger population of PwCS than our prior experiments, with participants exhibiting a wide range of gait function. While analyzing the study's data, several participants exhibited atypical gait patterns that caused our original definitions of the start and end of steps to produce results that did not match the actual gait behavior. Examples of these gait patterns are illustrated in the top row of Figure B1.

Some participants exhibited occasional instances in which their heel alternated anterior and posterior motions while their contralateral heel continued to move posteriorly (Fig. B1A). This pattern occurred when participants experienced a "stumble-like" event, or when their first attempt to advance their swing leg failed in placing the swing foot in front of the body. Our original method of identifying step start and end events included this entire period within a single step, which does not accurately represent the stepping behavior. A variation of this pattern was observed in a single participant, as illustrated in Figure B1B. This participant walked with a unique pattern in which they rotated their trailing paretic leg around a vertical axis through the ball of their foot just prior to taking a step with this leg (perhaps most easily visualized by "squishing a bug" beneath the ball of the foot). This pattern caused the velocity of the paretic heel to briefly become positive before the leg began swinging forward, leading to the start of paretic steps being detected falsely early. Finally, some participants occasionally spent a very long time in double support, often preceding paretic steps (Fig. B1C). Speculatively, this may have been due to participants ensuring they were confident in their balance before beginning the step. These long double support phases caused our original method of identifying step start and end events to report very long duration steps, not reflective of the actual stepping behavior.

Based on the above observations, we revised our analyses used to define the start and end of steps, with the goal of more accurately reflecting the stepping behaviors used to ensure mediolateral walking balance. To do so, we changed our analyses to be based on the anteroposterior velocity of the entire foot segment, rather than just the heel. Marker position data were filtered using a 4$^{th}$ order Savitzky-Golay smoothing filter with a 7-sample window. Any gaps in visible marker trajectories of up to 20 samples were filled using a cubic spline fit. The foot segment's center of mass position was then estimated from visible markers on the heel, second toe, first metatarsal, fifth metatarsal, and on a 3-marker rigid cluster secured over the dorsum of the foot. The goal of this increased complexity was to increase measurement reliability, minimizing the negative impact on event detection if the heel marker signal was compromised (e.g., due to marker occlusion).

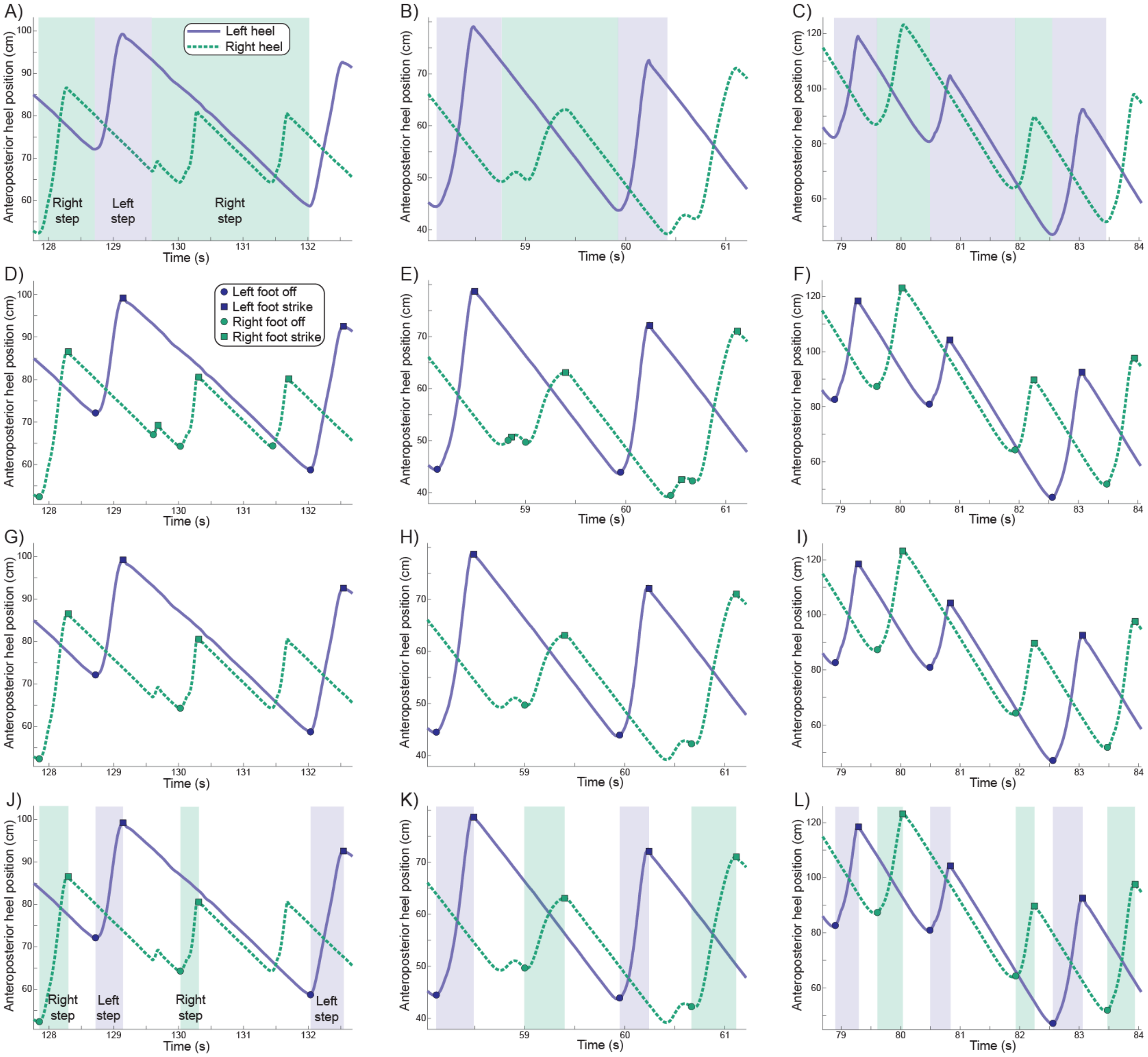


**Figure B1.** The progression of our revised method to identify the start and end of steps is illustrated as one moves down the figure. Each column illustrates an example of a distinct gait pattern. The top row (A-C) illustrates our original method for identifying steps, with the shaded areas indicating all complete steps during the illustrated period. The second row (D-F) illustrates the foot off and foot strike events identified using the estimated foot center of mass. While the event timing is quite similar to the events detected based just on heel velocity, small shifts are present – particularly for the false gait events due to foot rotation in panel E. The third row (G-I) illustrates only the foot off and foot strike events that corresponded to the largest anterior progression. In the fourth row (J-L), the shaded areas indicate all complete steps during the illustrated period, using our new method. The step duration is substantially reduced compared to our original method and is also less variable when unusual step patterns occur.

We then identified all timepoints at which the foot velocity changed anteroposterior direction for both legs. Timepoints at which the velocity changed from posterior to anterior were termed “foot off” (as the term “toe off” does not reflect the behavior seen in a subset of PwCS), and timepoints at which the velocity changed from anterior to posterior were termed “foot strike” (as the term “heel strike” does not reflect the behavior in a subset of PwCS). These gait events are illustrated in the second row of Figure B1.

We subsequently assumed that normal forward progression must follow a stereotypical sequence of gait events (e.g., right foot strike; left foot off, left foot strike, right foot off; right foot strike; etc.). All gait events that followed this sequence were accepted as being accurate representations of the gait pattern. Deviations from the standard sequence (as was the case in panels A and B of Figure B1) were interpreted as not being reflective of an individual’s steady-state stepping behavior. In this case, our goal was to determine which of the alternating periods of anterior and posterior velocities were most reflective of taking a forward step. Therefore, of the candidate steps, we identified the one in which the swing foot advanced the farthest anterior (accounting for the backward velocity of the treadmill belt). The results are shown in the third row of Figure B1.

Finally, the very long double support phases observed in some steps by some participants (see Fig. B1C) suggested that our previous definition for the end of a step did not constrain our analyses to the stepping behavior itself, the focus of the present study. An alternative approach is to define the end of the step as the timepoint when the stepping foot’s velocity changes from anterior to posterior [7]. Therefore, we compared the variability in calculated step duration using these two methods. For each leg, we calculated the standard deviation in step duration across all steps taken in the self-selected condition. We found that the step duration was significantly ($p<0.0001$) less variable when the end of the step was defined as the stepping foot starting to move backward (0.036±0.018 s for paretic steps; 0.029±0.013 s for non-paretic steps) than when the end of the step was defined as the contralateral foot starting to move forward (0.068±0.046 s for paretic steps; 0.073±0.060 s for non-paretic steps). We thus chose to define the end of steps as occurring when the stepping foot starts to move backward, as illustrated in the bottom row of Figure B1.

### Appendix B References


[1] K.H. Stimpson, L.N. Heitkamp, J.S. Horne, J.C. Dean, Effects of walking speed on the step-by-step control of step width, J. Biomech. 68 (2018) 78–83.

[2] K.H. Stimpson, L.N. Heitkamp, A.E. Embry, J.C. Dean, Post-stroke deficits in the step-by-step control of paretic step width, Gait Posture. 70 (2019) 136–140.

[3] L.N. Heitkamp, K.H. Stimpson, J.C. Dean, Application of a Novel Force-Field to Manipulate the Relationship Between Pelvis Motion and Step Width in Human Walking, IEEE Trans. Neural Syst. Rehabil. Eng. Publ. IEEE Eng. Med. Biol. Soc. 27 (2019) 2051–2058.

[4] N.K. Reimold, H.A. Knapp, R.E. Henderson, L. Wilson, A.N. Chesnutt, J.C. Dean, Altered active control of step width in response to mediolateral leg perturbations while walking, Sci. Rep. 10 (2020) 12197.

[5] N.K. Reimold, H.A. Knapp, A.N. Chesnutt, A. Agne, J.C. Dean, Effects of targeted assistance and perturbations on the relationship between pelvis motion and step width in people with chronic stroke, IEEE Trans. Neural Syst. Rehabil. Eng. Publ. IEEE Eng. Med. Biol. Soc. PP (2020).

[6] H.A. Knapp, B.A. Sobolewski, J.C. Dean, Augmented Hip Proprioception Influences Mediolateral Foot Placement During Walking, IEEE Trans. Neural Syst. Rehabil. Eng. Publ. IEEE Eng. Med. Biol. Soc. 29 (2021) 2017–2026.

[7] J.A. Zeni, J.G. Richards, J.S. Higginson, Two simple methods for determining gait events during treadmill and overground walking using kinematic data, Gait Posture. 27 (2008) 710–714.
