## Appendix C for "Relationships between mediolateral step modulation and clinical balance measures in people with chronic stroke"

The biomechanical gait metrics presented in this manuscript's main text focused on the relationship between pelvis dynamics at the start of the step and the body's mediolateral configuration at the end of the step. The reason for this focus is because in order to execute active control over the stepping behavior based on pelvis dynamics, such dynamics must be sensed early in a step, not just at the end. An alternative approach used by other groups focuses on the relationship between pelvis dynamics at the end of the step and the body's mediolateral configuration at the same time point. We here present results based on that time point.

As with our primary results, the strength of all detected significant relationships was only weak to moderate. Overall, the form of the observed relationships were similar to those focused on step start pelvis dynamics, although less likely to reach statistical significance. For steps taken with both the paretic and non-paretic leg, higher $\rho_{FP}$ values were linked with better FGA scores and less fear of falling. Higher paretic $\rho_{SW}$ values were also linked with better FGA scores. No biomechanical gait metrics exhibited a significant relationship with either ABC score or fall history.

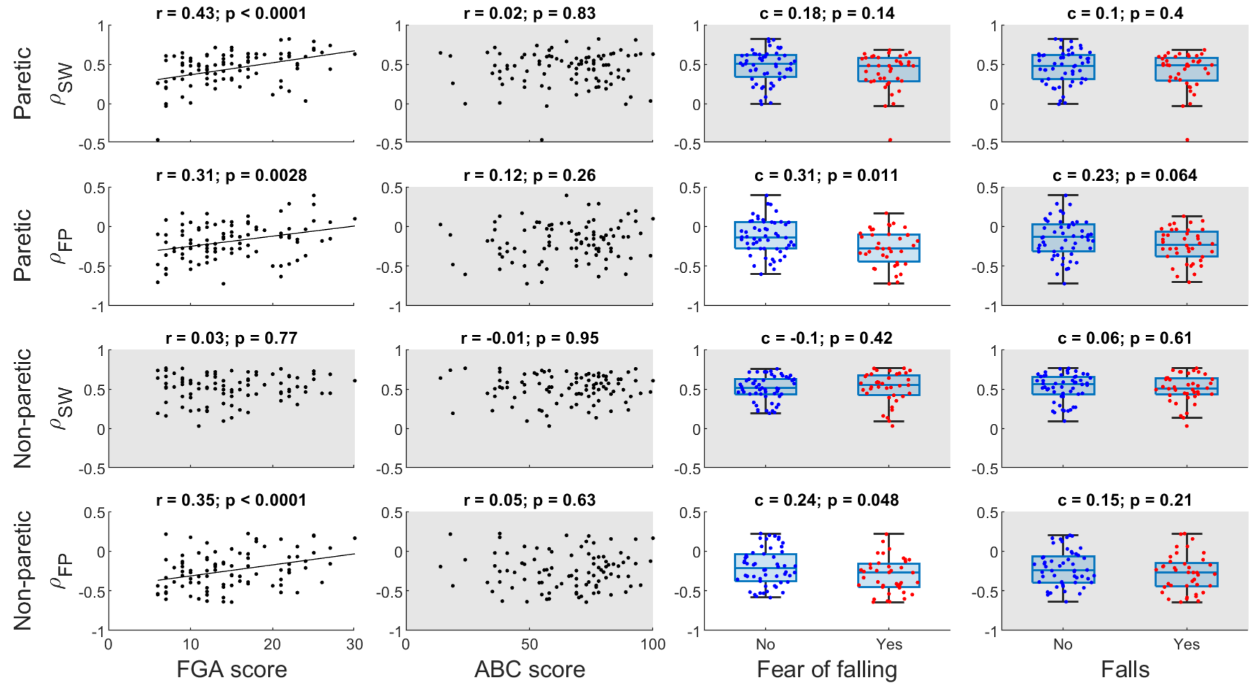


**Figure C1.** This figure follows the same structure as Figure 2 from the main text, but presents partial correlations calculated based on pelvis displacement at the end of the step, rather than the start. For all panels, dots represent individual data points, and effect sizes and p-values are presented at the top. Shaded panels indicate that no significant relationship was detected. Relationships for $\rho_{PD}$ are not presented, as this metric trivially converges to 1 at the end of a step.
